# The nanos^d^ integral gene drive enables population modification of the malaria vector *Anopheles gambiae*

**DOI:** 10.1101/2025.05.02.651860

**Authors:** Pei-Shi Yen, Sebald ANR Verkuijl, Paolo Capriotti, Giuseppe Del Corsano, Astrid Hoermann, Maria Grazia Inghilterra, Irati Aramburu-Gonzalez, Moeez A Khan, Dina Vlachou, George K Christophides, Nikolai Windbichler

**Author notes:** These authors contributed equally (Pei-Shi Yen, Sebald ANR Verkuijl, Paolo Capriotti, Giuseppe Del Corsano).

## Abstract

The modification of mosquito populations at scale through CRISPR-Cas9-mediated homing gene drives is a promising route for malaria vector control. Integral gene drives (IGDs) are designed to utilize the regulatory sequences of endogenous genes to express only the minimal set of components required for gene drive. In this study, we describe the creation and characterization of the nanos^d^ IGD targeting and inserted into the *nanos* gene of the malaria vector *Anopheles gambiae* and show that it achieves high rates of gene drive (97.7% in females, 99.0% in males). We find that homozygous nanos^d^ females but not males show impaired fecundity, and a variable loss of ovary phenotype. Transcriptomic analysis of ovaries points to decreased transcript levels of the *nanos* gene when harbouring Cas9. As a minimal genetic modification, nanos^d^ does not induce widespread transcriptomic perturbations, and its susceptibility to *Plasmodium* spp. and O’nyong nyong virus infection remains similar to wild-type mosquitoes. Importantly, we find that nanos^d^ propagates efficiently in caged mosquito populations and is maintained as a source of Cas9 after the emergence of drive resistant alleles whilst also mobilising a non-autonomous antiparasitic effector modification. The nanos^d^ gene drive shows promise as a genetic tool for malaria vector control via population modification, and we outline steps towards its further optimization.

## Introduction

Malaria is the deadliest infectious disease transmitted by mosquitoes. In 2023, an estimated 263 million malaria infections occurred, leading to 597,000 deaths^1^. This follows a trend of increased infection and death incidences in recent years, driven by, among others, the spread of drug, diagnostic, and insecticide resistance. Additionally, climate change is altering vector distribution and influencing the malaria transmission cycle^2^, while the lingering healthcare disruptions from the COVID-19 pandemic have further weakened prevention and treatment efforts^1,3^. This underpins the need for alternative tools and strategies for malaria control, particularly methods that target mosquito vectors and reduce transmission. Among the proposed strategies for vector control, homing gene drive systems are considered one of the most promising tools to achieve malaria eradication^4^. Gene drives are genetic elements engineered to bias their inheritance, facilitating the spread of specific traits within a target population. Gene drive systems can allow traits of interest to increase in frequency across generations, even in the absence of a fitness advantage. The advancement of the CRISPR/Cas9 endonuclease platform has significantly accelerated the development of gene drive technologies for public health and pest control applications^5^ In a homing gene drive system, the CRISPR/Cas9 nuclease induces a DNA double-strand break at a specific locus. The cellular repair machinery then uses homologous sequences from the other chromosome that contains the gene drive components. This ‘homing’ mechanism converts the target sequence into a gene drive carrying allele in the germline thus increasing drive inheritance. Several homing drives have demonstrated high levels of biased inheritance in *Anopheles* mosquitoes^6–11^. Homing drives typically consist of a Cas9 coding sequence driven by germline-specific regulatory elements, and a gRNA expressed under the control of an RNA Polymerase III promoter. The fitness costs, homing efficiency, and resistance rate of the gene drive is strongly influenced by the nuclease activity window. Activity in non-germline tissues, through ‘leaky’ expression or parental nuclease deposition into the zygote, can be detrimental. When they target important mosquito genes, gene drives are designed to ensure that the nuclease activity window (germline) does not overlap with that of the target gene (somatic). This is to make use of the favourable DNA repair preference in the germline, ensure that recessive somatic fitness cost phenotypes do not manifest in heterozygote drive carriers^9,12^, and prevent non-specific stress from DNA damage in tissues that do not contribute genetic material to the gametes. However, it is challenging to achieve exclusively germline expression, and gene drive optimisation efforts generally focus on testing a range of putative regulatory elements from germline-specific genes to drive Cas9 expression^13–15^. This optimisation approach has had only limited success, and suboptimal performance is commonly attributed to not capturing all the regulatory elements specific to the germline gene’s expression (often only a promoter fragment, 5‘UTR, and 3‘UTR), and interference from cis-regulatory elements at the transgene’s integration site within the genome (e.g., enhancers)^9,12^.

In contrast to conventional homing gene drives, integral gene drives (IGDs) seek to utilise the full regulatory context of an endogenous germline gene locus, making it more likely that the intended expression pattern is achieved. They do this by being inserted in-frame with and maintaining the function of the endogenous expression unit of the germline gene. For integral drive elements, there are no intended recessive fitness costs of the insertion, and as such activity of the nuclease can overlap with that of the target gene in the germline. IGD benefit from a reduced genetic footprint size and make it more likely that mutations that disrupt the drive components are selected against though loss of function of the host gene. IGDs have been demonstrated in *Drosophila melanogaster, Mus musculus, Anopheles stephensi*, and *Anopheles gambiae*, and while allowing for efficient homing rate, they can incur severe unintended fitness costs^16–19^. Additional efforts are required to identify suitable host gene loci and gene drive cassette designs that allow for optimal IGD performance.

Regulatory elements from germline genes that have worked sub-optimally in traditional gene drive designs for the reasons outlined above may work well in an IDG design^20^. The *nanos* gene is part of a highly conserved family found in both vertebrates and invertebrates^21,22^. In *A. gambiae, nanos* expression is restricted to the male and female germline as well as embryos^23,24^, where it encodes a zinc-finger RNA-binding domain involved in regulating the translation of mRNAs critical for polarity and germ cell development in embryos^25,26^. Regulatory sequences (including promoter sequences and UTRs) of the *nanos* gene have been utilised in gene drives and other genetic modifications targeting the germline of *Anopheles* species^27–30^. Here, we utilise the *nanos* locus of *A. gambiae* for the integration of an IGD construct designed to express Cas9 protein under the control of *nanos* regulatory elements, along with a *nanos*-targeting gRNA expression cassette embedded within a synthetic intron, the *nanos* IGD can trigger efficient homing in the germline. To achieve population modification, the *nanos* IGD is designed to spread naturally through populations without intended fitness costs. In combination with a non-autonomous anti-malarial effector, this drive aims to disseminate itself and the effector within the target population. To ensure utility in anti-malaria efficacy, the genetic modification in *nanos* IGD mosquitoes should not affect vector competence for malaria parasites or other relevant human pathogens. The goal is to evaluate the feasibility and effectiveness of this gene drive and to assess its potential for sustainable disease control.

## Results

An integral gene drive construct targeting *A. gambiae* was engineered for insertion into the *nanos* locus (AGAP006098) via CRISPR/Cas9-mediated homologous recombination (**Fig. 1a**). The construct includes an NLS-Cas9-NLS coding sequence with an E2A skipping peptide facilitating the in-frame fusion with the endogenous *nanos* coding sequence and flanked by *nanos* homology arms. An artificial intron that contains a gRNA and GFP fluorescent marker expression cassette was embedded within the E2A skipping peptide sequence. Upon embryonic microinjection, Cas9 provided by a helper plasmid and *nanos*-targeting gRNA transcribed from the donor plasmid mediate HDR at the target site. A gene drive strain carrying a GFP selection marker (nanos^D^) was generated through crossing the GFP-positive G_0_ individuals with wild-type mosquitoes, and a marker-free strain (nanos^d^) was subsequently established by outcrossing to a transgenic mosquito line expressing Cre recombinase to excise the selection marker flanked by loxP sites.

**Figure 1.**
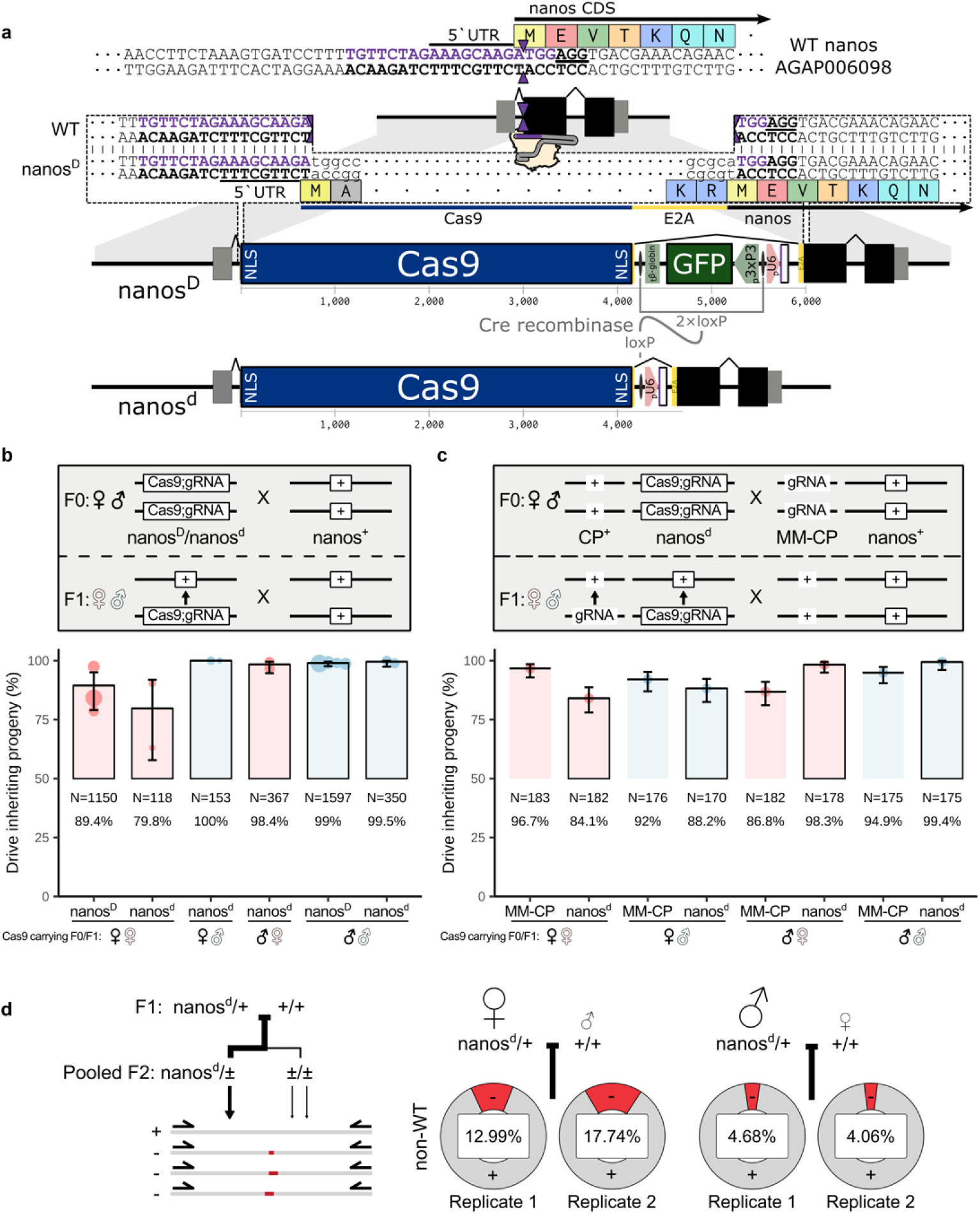
The *nanos* integral gene drive. **(a)** Schematics represent the *nanos* integral gene drive. The sequence of the *nanos* locus, gRNA target site (sequence in purple), PAM site (underlined), and CRISPR/Cas9 cutting site (purple triangles) are highlighted. To generate nanos^D^, the *nanos* IGD construct was integrated immediately upstream of the *nanos* CDS via homologous recombination. The marker-free nanos^d^ line was established by crossing nanos^D^ mosquitoes to a *Cre* recombinase line. **(b)** Drive transmission from heterozygous male or female (coloured gender symbols) nanos^D^ (D+) and nanos^d^ (d+) individuals. **(c)** Drive transmission from a double heterozygous male or female nanos^d^;MM-CP individuals. Sex of the nanos^d^ grandparent is indicated with black gender symbols (contribution of MM-CP is from the opposite sex). Each point represents the mean from a pooled independent biological replicate, with the GLM estimated mean and the total number of progeny scored listed. Bars start at the expected inheritance rate and extend to the observed overall mean inheritance. **(d)** The proportion of indels identified by amplicon sequencing analysis of the offspring of male and female nanos^d^ heterozygote crosses with the wild-type mosquitoes.

Highly efficient inheritance rates (79.8–100%) were observed for the nanos^D^ and markerless nanos^d^ drive lines via fluorescent and PCR-based screening, respectively (**Fig. 1b**). Notably, crosses with the possibility for maternal deposition into female drive heterozygotes performed worse (inheritance rates of individuals from a female parent and grandparent (♀♀): 79.8% and 89.4% for nanos^d^ and nanos^D^, respectively) than those with paternal nanos^d^ contribution (♂♀/♂♂; 98.4–99.0%) or those with maternal contribution to male drive heterozygotes (100%). This suggests an impact of maternal deposition on drive efficiency only in female drive carriers. A reduction in the observed inheritance bias in the female crosses (derived from female grandparents; ♀♀) after removal of the marker may indicate the reporter expression cassette within the artificial intron does not necessarily affect the homing rate (*p* = 0.29).

We assessed the ability of nanos^d^ to simultaneously induce homing of a non-autonomous anti-malarial effector (MM-CP) that carries a gRNA targeting the *zinc carboxypeptidase A1* gene^31^. Male and female nanos^d^;MM-CP double-heterozygous individuals were crossed with wild-type mosquitoes, leading to efficient inheritance rates of 84.1–99.4% and 86.8–96.71% for the nanos^d^ and MM-CP elements, respectively (**Fig. 1c**). While overall inheritance rates did not significantly differ between the two loci (*p* = 0.069), inheritance was significantly higher in the crosses involved a male Cas9 or gRNA grandparent (*p* = 0.001 and *p* < 0.0001, respectively). The nanos^d^ inheritance rates were higher than MM-CP in the crosses with male nanos^d^ grandparents (♂♀/♂♂), while the MM-CP inheritance rates were higher in the crosses with female nanos^d^ grandparents (♀♀/♀♂). This would be consistent with the effect of maternal deposition being more substantial for nanos^d^, as both gRNA and Cas9 are present in the grandmother, while the MM-CP gRNA element was always contributed by the grandparent lacking Cas9. Crosses with a GFP-marked CP element and nanos^d^ showed similar inheritance rates to those with MM-CP (**Fig. S1**).

To further investigate the impact of maternal deposition and the nature of the resistance alleles it generated, pools of nanos^d^-homozygous males (N = 40 and 49) and females (N = 20 and 40) were crossed with wild-type mosquitoes, and the non-nanos^d^ alleles, expected to include both unmodified as well as mutant alleles, were amplified and sequenced in approximately 400 L2-L3 larvae offspring. Insertions and deletions (indels) caused by non-homologous end joining (NHEJ) were identified within the *nanos* locus in 4.06% and 4.68% of non-nanos^d^ reads from nanos^d^ male parents, while 12.99% and 17.74% are from nanos^d^ female parents (**Fig. 1d**), suggesting additional resistance alleles generated due to maternal deposition. In line with this, the indel(s)-containing alleles recovered from the male cross have low diversity, suggesting F1 germline mutations, while these indel(s)-containing non-wildtype alleles in the female cross were more diverse, implying F2 mosaicism due to maternal deposition of Cas9 (**Fig. S2**). These indels were predicted to disrupt the Cas9 target site, forming resistance alleles against nanos^d^, potentially limiting the spread of the gene drive. Interestingly, while many of the observed mutations were predicted to abolish nanos expression a significant set of alleles from maternal nanos^d^ crosses reconstituted the nanos start codon (**Fig. S2**). This could indicate that in these experiments some level of active selection, at the level of the embryo, would help to maintain *nanos* function.

Next, the life history traits of the nanos^d^ gene drive were evaluated to identify potential unintended fitness costs. Fecundity was measured by counting the number of eggs laid by individual inseminated females with different nanos^d^ zygosity crossed to wild type mosquitoes (**Fig. 2a**). No significant differences in egg counts were observed in crosses between nanos^d^ homozygous males and wild type females, or between intercrossed nanos^d^ heterozygous males and females (male nanos^d^ grandparent). However, nanos^d^ homozygous females mated with wild-type males produced significantly fewer eggs compared to wild-type females, an effect that was larger in nanos^D^ mosquitoes (**Fig. S3**). Aside from reduced fecundity, no significant differences were observed for fertility (**Fig. 2b**), larval survival (**Fig. 2c**), or the sex ratio (**Fig. 2d**) between nanos^d^ and wild-type mosquitoes. The nanos^d^ mosquitoes exhibited increased time to pupation (**Fig. 2e**) and a somewhat reduced adult lifespan (**Fig. 2f**).

**Figure 2.**
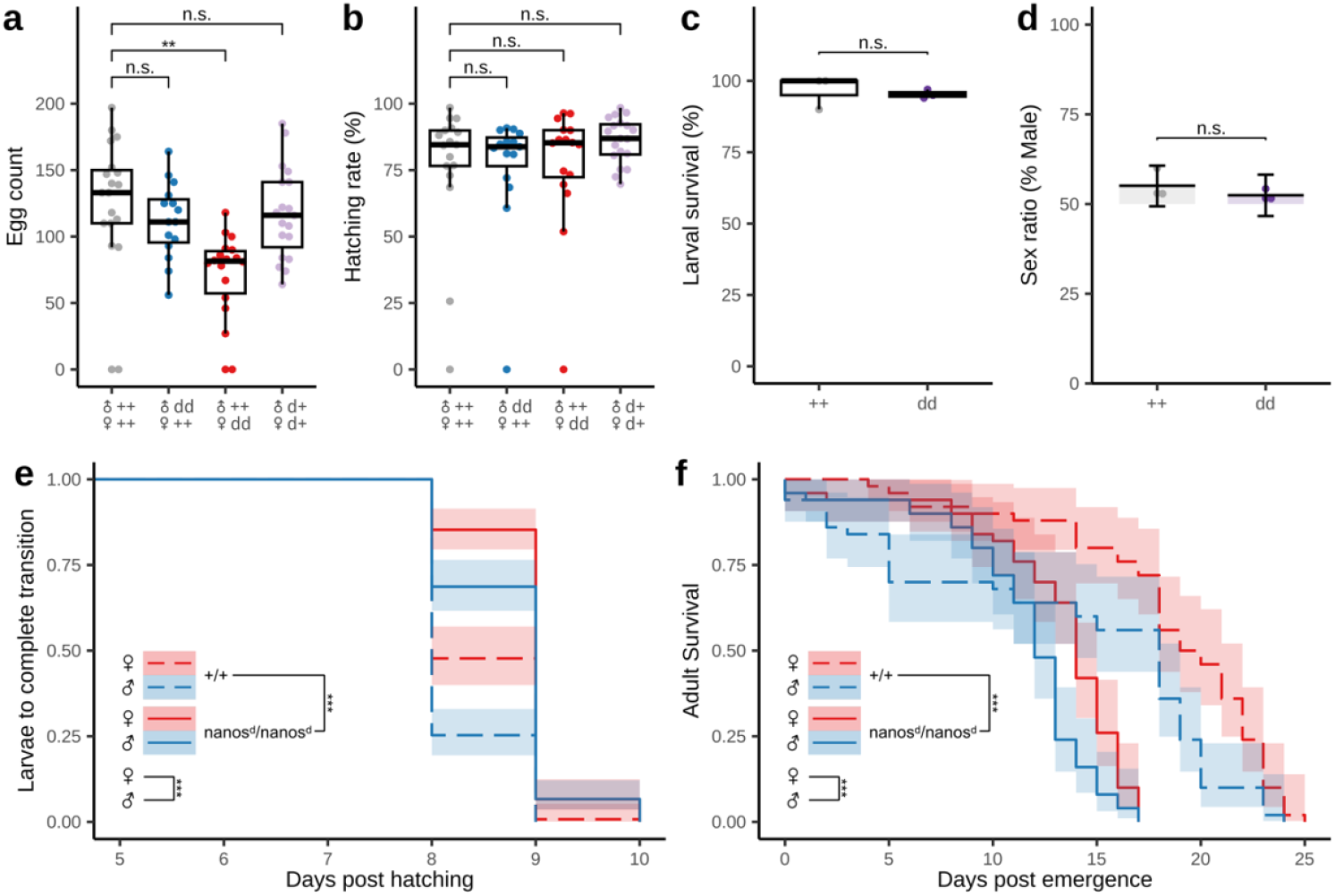
Fitness and life history traits of nanos^d^ mosquitoes. **(a)** Fecundity and **(b)** fertility of females with different nanos^d^ zygosity, or crossed to nanos^d^ carrying males. The heterozygous nanos^d^ mosquitoes in the fecundity and fertility tests were the progeny of male nanos^d^ crossed to female wild-type mosquitoes. Egg count significance levels were calculated using an unpaired two-tailed Student’s *t*-test using Bonferroni correction. **(c)** Larval survival, **(d)** pupal sex ratio, and **(e)** development time from larval to pupal stages for homozygous nanos^d^ individuals. For b-d, significance levels were calculated using a binomial GLM with replicate as a random effect and Dunnett’s multiple comparisons test. **(f)** Averaged daily survival of homozygous male or female nanos^d^. The time series analysis in e and f was conducted using a mixed-effects Cox proportional hazards model. Pairwise comparisons were averaged over the effect of sex, using Tukey’s method with *P*-value adjustment. Shaded areas indicate the 95% pointwise confidence intervals.

The reduced fecundity phenotype was further investigated by dissecting and examining mosquito ovaries (**Fig. 3a**). Results showed that 53.5% of sugar-fed nanos^d^ mosquitoes had fully or partially underdeveloped ovaries, whereas only 5.71% of wild-type mosquitoes exhibited the same phenotype (**Fig. 3b**). The morphology of the nanos^d^ ovaries was generally less compact compared to fully developed ovaries from wild-type mosquitoes.

**Figure 3.**
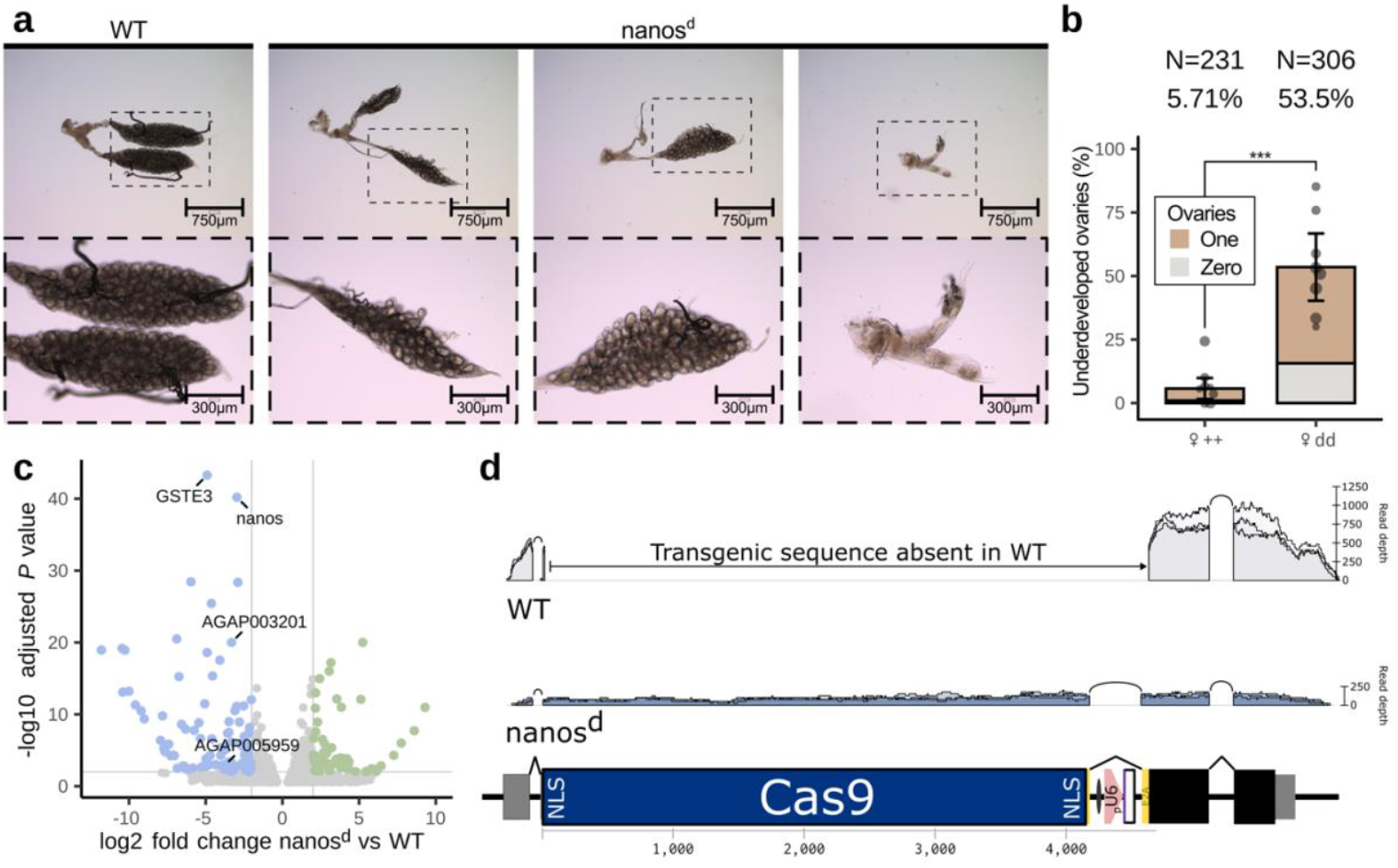
Development and gene expression profile of nanos^d^ mosquitoes’ ovaries. **(a)** Ovarian morphology of wild-type and nanos^d^ mosquitoes, indicating severe developmental defects in nanos^d^ ovaries, which appear fully or partially underdeveloped and more dispersed. **(b)** Number of underdeveloped ovaries in nanos^d^ and wild-type mosquitoes. Mean estimates and significance levels were calculated using a multinomial GLM with replicate as a random effect. **(c)** Volcano plot of differentially expressed genes in nanos^d^ and wild-type ovaries of 2– 4-day-old mosquitoes, significant transcriptomic changes were highlighted, including genes involved in germline development. Differentially expressed genes between nanos^d^ and wild-type mosquitoes, identified based on a significance threshold of *P* ≤ 0.01 and a log_2_-fold change ≥ 2, are highlighted. **(d)** Reads mapped to *nanos* locus and the nanos^d^ construct, demonstrating Cas9 expression and confirming correct Cas9 gene splicing in the ovaries of nanos^d^ mosquitoes. The total number of reads mapped to the genome for the two groups was similar (Ifakara: 4.2 ’ 10^7^; nanos^d^: 4.1 ’ 10^7^).

We next performed transcriptome analysis on the ovaries from nanos^d^ and wild-type females (**Fig. 3c**). Gene expression profiling revealed a significant reduction in *nanos* expression levels, with only one other gene the glutathione S-transferase gene, *GSTe3* (AGAP009197), typically upregulated in DDT-resistant strains^32^, reaching a similar level of significance (**Fig. 3c,d**). We confirmed that the additional intron we introduced as part of the nanos^d^ construct did not lead to unintended splicing products (**Fig. 3d**). GO enrichment analysis indicated that the expression levels of several genes involved in both female and male dipteran fertility were affected. Some female fecundity-related genes were significantly downregulated in the ovaries of nanos^d^ mosquitoes and may be linked to the reduced fecundity observed in this study. The maternally-expressed AGAP003201 is the ortholog of *Drosophila Tao-1* gene, which is involved in cell migration during embryogenesis in *Drosophila*^33^; *yellow-g2* (AGAP005959) was also found to be downregulated in the ovaries of nanos^d^ mosquitoes. In *Aedes albopictus*, ovary-expressed *yellow-g2* encodes a dopachrome-conversion enzyme essential for cuticle melanisation, which is required for egg desiccation resistance and pigmentation^34^.

Infection experiments were conducted to assess the susceptibility of nanos^d^ mosquitoes for pathogen infection (**Fig. 4a**). The results showed that female nanos^d^ mosquitoes do not exhibit an increased infection rate (prevalence) for the rodent malaria parasite *P. berghei*, or for the human parasite *P. falciparum* (**Fig. 4b**). In fact, the *P. falciparum* oocyst numbers were slightly lower in nanos^d^ mosquitoes compared to wild type controls (**Fig. 4c**), accompanied by slightly larger oocysts likely due to reduced competition for resources among parasites (**Fig. 4d**). We also assessed vector competence for O’nyong nyong virus (ONNV). Again, nanos^d^ mosquitoes exhibited no increase in infection rates (**Fig. 4e**), or viral replication (**Fig. 4f)**. The absence of detectable viral particles in the heads and saliva of infected mosquitoes suggests a low risk of ONNV dissemination and transmission, indicating that the genetic modification does not enhance virus replication.

**Figure 4.**
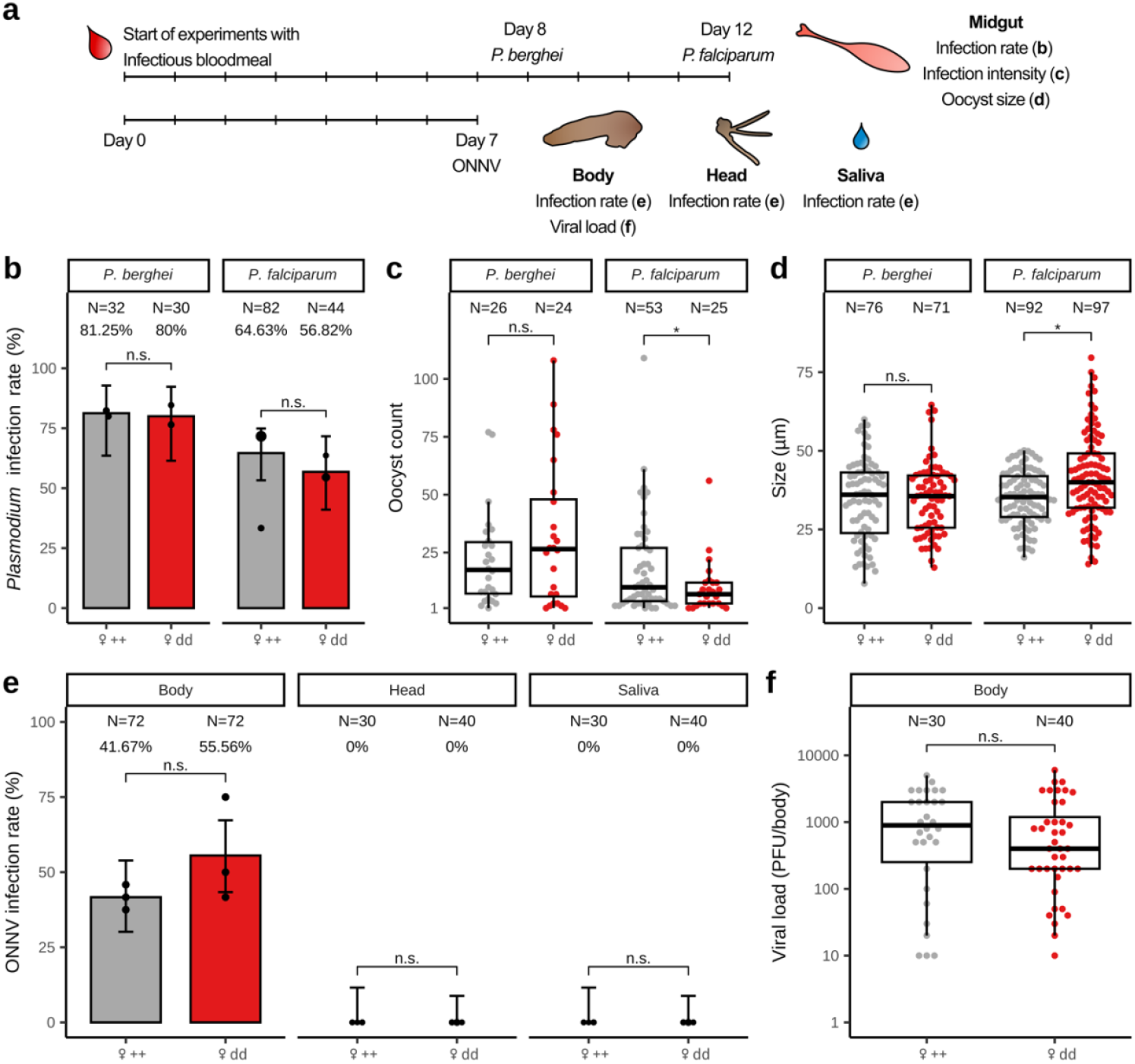
Permissiveness to pathogen infection. **(a)** Timelines for *Plasmodium* spp. (upper) and ONNV infection experiments (bottom). **(b)** *P. falciparum* and *P. berghei* oocyst infection rate, **(c)** infection intensity **(d)** and oocyst size in the midgut of wild type and nanos^d^ mosquitoes at 8 or 12 days post-blood feed (dpbf). Pooled results of two replicates are presented. The median oocyst number in each midgut is indicated. **(e)** ONNV infection rate in mosquito bodies, heads, and saliva at 7 dpbf. **(f)** Viral load comparison between nanos^d^ and wild-type midguts. Means and significance levels for b and e were calculated using a binomial GLM with replicate as a random effect. Significance levels for c, d, and f were calculated using an unpaired two-tailed Student’s *t*-test using Bonferroni correction.

We next assessed the capacity of nanos^d^ to drive its own propagation and that of the non-autonomous effector MM-CP in four independent cage populations. Each cage was seeded with 400 individuals, consisting of 200 nanos^d^ heterozygotes and 200 MM-CP heterozygotes at a 1:1 sex ratio. This corresponds to an initial transgene allele frequency of 25% (50% carrier frequency) for both nanos^d^ and MM-CP. Transgene carrier (**Fig. 5a** and **Fig. S4**) and allele (**Fig. 5b** and **Fig. S5**) frequencies were monitored over 20 discrete generations using PCR genotyping, as neither line carries a fluorescent marker.

**Figure 5.**
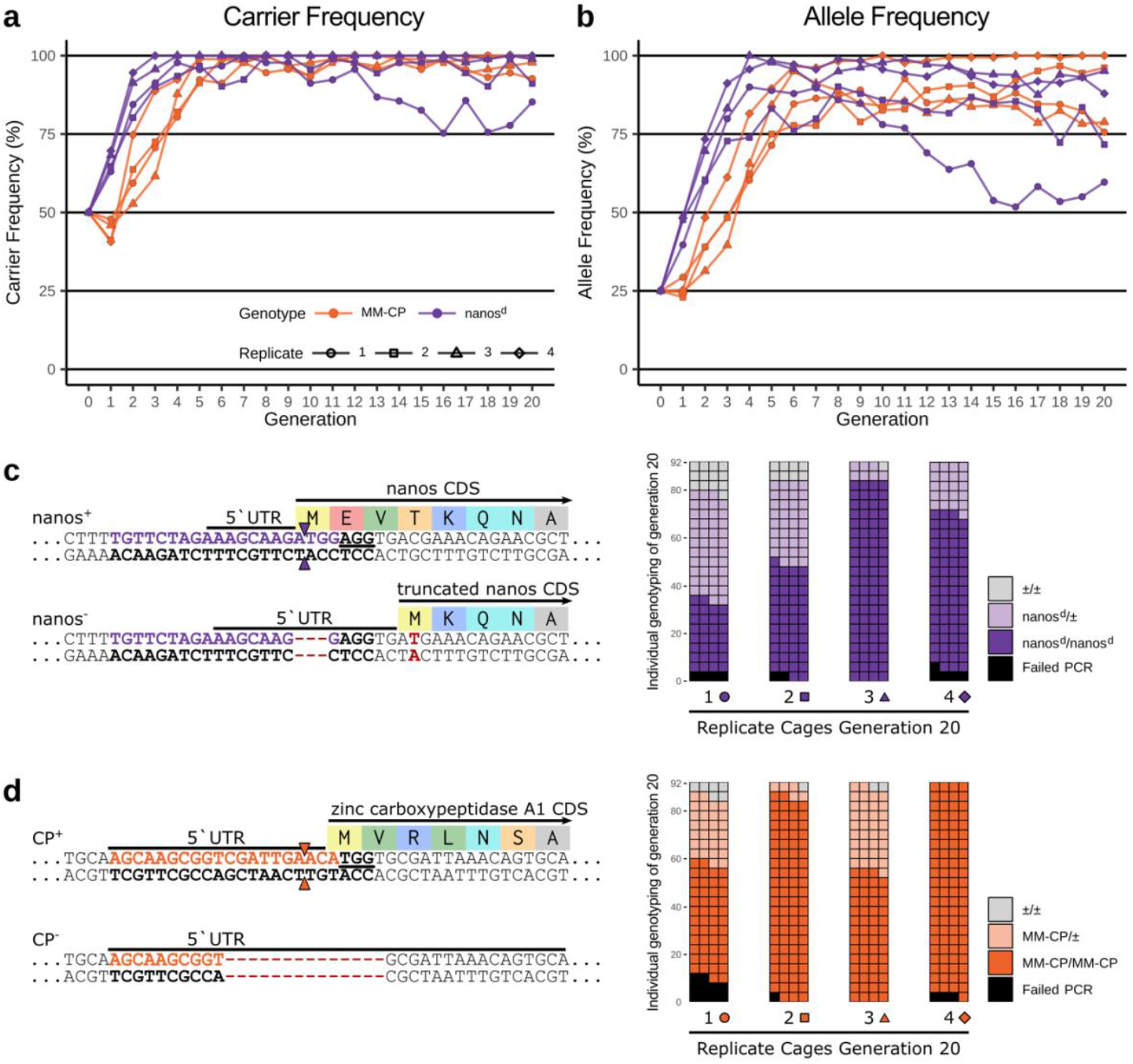
Population convention experiments under laboratory conditions. **(a)** Carrier and **(b)** allele frequency over 20 generations nanos^d^ (purple) and MM-CP (orange), in four independent trials. Populations were seeded with 50% nanos^d^ and 50% MM-CP heterozygotes. The number of MM-CP and nanos^d^ transgene frequencies and carriers was determined across multiple generations by molecular genotyping. **(c)** Reference sequences for *nanos* and *zinc carboxypeptidase A1* are shown, with the PAM highlighted with a black underline. The gRNA target sites are indicated by solid arrows. The predominant resistance alleles exhibit deletions around the cleavage site, leading to sequence modifications. In the *zinc carboxypeptidase A1* locus, the resistance allele features a deletion in the target sequence, which prevents Cas9-mediated cleavage. In nanos^d^ mosquitoes, a deletion near the PAM site results in the shifting to an alternative start codon (highlighted in scarlet). The waffle plots represent genotype composition of MM-CP and nanos^d^ in each cage at the 20^th^ generation.

Both transgenes spread rapidly, reaching a plateau after 5-10 generations, with nearly all individuals carrying at least one copy of the transgenes. As expected, MM-CP trailed behind nanos^d^ in frequency, as its biased inheritance depends on the presence of nanos^d^ in the same individual, which could not occur in the initial release generation. In the first replicate, the nanos^d^ frequency began to decline after generation 7, suggesting the emergence of functional drive-resistant alleles. The subsequent stabilization of the nanos^d^ frequency and the lack of rapid loss suggests that any associated fitness cost is primarily recessive.

To characterize potentially functional drive-resistant alleles in these cage populations, individuals that lacked nanos^d^ at the *nanos* locus were sampled and sequenced. The results revealed a prominent allele with a 3bp deletion at the start codon of the *nanos* gene (**Fig. 5c**), which was also prominent in sequencing of pooled crosses (**Fig.S2**). While this mutation disrupts the *nanos* coding sequence, it was found in combination with a downstream C to T substitution that introduces an alternative start codon (Thr3Met), resulting in three-amino acid N-terminal truncation. This variant may preserve *nanos* function and thus confer resistance to the drive without a major fitness cost. Supporting this, RNA sequencing data confirmed that Thr3Met is common variant in our wild-type colony, suggesting it may represent pre-existing genetic variation, and that resistance could be circumvented with alternative gRNA design. In contrast, a 14bp deletion identified in the MM-CP cassette is predicted to disrupt the CP function entirely (**Fig. 5d**) Given that MM-CP frequencies remained high even when nanos^d^ frequencies declined, this mutation is unlikely to confer a selective advantage, consistent with a non-functional allele lacking fitness benefit.

## Discussion

Significant efforts are required to reach technical, social, and regulatory consensus on field testing and eventual deployment of gene drives. The separation of autonomous drive and non-autonomous effector transgenes allows for the independent development of such gene drive traits that have differing requirements related to their respective functions. The autonomous drive element requires faithful expression in the germline. Here, the integral gene drive paradigm offers to closely recapitulate germline expression of endogenous and minimise the genetic footprint of the autonomous gene drive modification. On the other hand, effector modifications targeting the parasite such as the MM-CP effector expressing anti-microbial peptides must be active in non-germline tissues, and as such must be housed at distinct loci with appropriate expression patterns. Thus, separating the effector and inheritance bias modules is an obvious step and comes with the benefit of phased testing approaches, which can provide a more gradual path to approval. The linkage of the anti-malaria or drive components to an endogenous gene could also increase transgene integrity by selection against loss of function mutations that also affect the host gene (*e*.*g*., frame shift mutations). We have utilised the *nanos* locus of *A. gambiae* to develop an autonomous integral gene drive element. This nanos^d^ gene drive element makes use of the regulatory regime of the *nanos* host gene and supports for high levels of inheritance bias. Despite the removal of the intronic fluorescent marker cassette, some fitness parameters, including fecundity, pupal developmental time, and adult lifespan, were affected in homozygous nanos^d^ individuals. The impact on fertility and effects on ovarian development we observed are likely a direct result of reduced *nanos* expression. While further optimization is therefore possible nanos^d^ nevertheless represents a marked improvement over previous attempts at the *zpg* locus and unlike these earlier designs was predicted to be a fully invasive gene drive element.

We noted a decrease in inheritance bias in female, but not male, nanos^d^ heterozygotes who were generated through crosses that allowed for maternal deposition of Cas9. Maternal deposition of Cas9, or Cas9;gRNA, into the zygote can lead to cutting at stages when homing is not favoured, and has been commonly observed for canonical *Anopheles* nanos drives^9,10,35^. Deposition can cause stochastic mosaic mutation patterns in germline and non-germline cell lineages alike. Crosses with male nanos^d^ and female MM-CP grandparents showed a lower MM-CP inheritance rate compared to nanos^d^. In contrast, crosses with female nanos^d^ and male MM-CP grandparents exhibited a lower nanos^d^ inheritance rate compared to MM-CP. These results may suggest that maternal deposition of both Cas9 and gRNA simultaneously could contribute to reduce inheritance rates more.

When we sequenced the offspring of drive heterozygotes, many of the indels identified resulted in frameshift mutations that are predicted to disrupt correct protein translation. However, due to the cut-site overlapping the start codon of nanos, a range of indels that extended into the 5’UTR recreated an alternative start codon, possibly allowing for continued expression of functional *nanos* (**Fig. S2**). Notably, in the cage trial, we identified a separate mutation that restored the *nanos* reading frame through an alternative downstream methionine codon, present in a subset of our wild-type colony. Functional mutations would then plausibly explain why some cage experiments saw a decline in the nanos^d^ allele frequency. This accumulation may affect the persistence of nanos^d^ but does not necessarily substantially impact the spread of the antiparasitic payload construct if an intermediate carrier frequency of Cas9 is reached and maintained within the population (**Fig. S4–5**). Such maintenance of a Cas9 source could even allow for the release and propagation of a secondary non-autonomous effector in a population. Nonetheless, deposition of Cas9 by *nanos* is expected to also negatively affect the gRNA-expressing effector and should therefore be minimised. Overall, we describe a *nanos*-IGD with minor to moderate fitness costs that can disseminate both an autonomous gene drive and a non-autonomous antiparasitic payload construct. The nuclease profile of this strain can be further optimised by shifting the target site and decreasing the persistence of Cas9 into the embryo by reducing the protein stability.

In *Drosophila melanogaster*, the *nanos* gene is expressed in the early germarium, which maintains transcriptional quiescence and acts as an important determinant of posterior patterning during oogenesis^36,37^. The *nanos* gene itself was one of the most significantly downregulated genes in nanos^d^ ovaries. While mRNA transcription rate and stability may have been affected by the insertion, we cannot exclude that the insertion affects *nanos* in a different way, such as delayed translation, or loss of protein functionality due to insufficient separation at the 2A linker. In that case, the lower levels of *nanos* transcripts and other fecundity-related genes in the ovaries may be an indirect effect of the underdeveloped ovaries phenotype. It is notable that the female nanos^d^ heterozygotes did not display a similar fertility defect compared to homozygotes. In the tissues where *nanos* is expressed, so is Cas9; if homing occurs rapidly, we should observe similar impacts in heterozygotes and homozygotes. Moreover, in heterozygotes, the two *nanos* alleles will be engaged in DNA repair processes that could prevent expression at either allele. The unaffected fertility of heterozygotes could be due to the accumulation of wild-type nanos transcripts before the homing process is initiated. This would most likely be due to delayed translation but could also be explained by the initial absence of gRNA expression when Cas9 is first translated. Additionally, a grandparental effect might contribute. Homozygotes would have received reduced maternal deposition of *nanos* transcripts from homozygous mothers, while heterozygotes from wild-type mothers would have received normal levels, potentially diminishing the effect during early developmental processes.

Another significant fitness impact was observed in adult lifespan, with both male and female nanos^d^ homozygous mosquitoes exhibiting reduced survival rates compared to wild type. The underlying causes of the shortened lifespan in both sexes remain unclear. Given the low genetic diversity of transgenic strains, inbreeding is a likely explanation for some of the somatic phenotypes observed. Although Cas9 expression may place a general metabolic load on cells, it is expected to be restricted to the germline at specific developmental stages and would be expected to have minimal impact on overall mosquito lifespan. The ubiquitous expression of gRNA molecules could also account for some systemic fitness effects. The reduced lifespan of females, if attributable to the genetic modification itself, could be a desirable trait for reducing the risk of pathogen transmission. In general, *Anopheles* mosquitoes require 10-14 days to transmit sporozoites after ingesting human *Plasmodium* gametocytes^38^ and approximately 7 days for ONNV transmission.

Along similar lines the genetic modifications introduced at the *nanos* locus and a significant reduction of nanos transcript observed in the ovaries, are effects expected to be confined to the reproductive system of mosquitoes and would not be expected to altering immune responses against pathogen infections. Again, the ubiquitous gRNA expression could trigger more systemic changes to immunity. We found that nanos^d^ mosquitoes showed no increased vector competence for *Plasmodium* or ONNV, although their reduced adult lifespan would limit the chance for pathogen transmission and prevent the mosquito from becoming transmissible. In four long-term cage studies, we demonstrated that nanos^d^ could effectively propagate itself and the MM-CP non-autonomous effector modification. The creation and accumulation of resistance alleles within the target population remain challenges in optimizing *nanos*-IGDs for population control. Designing guide gRNA sequences that target essential motifs further removed from the start codon, is expected to mitigate the accumulation of the type of resistance alleles we observed. Moreover, CRISPR/Cas9 gene drives employing multiple gRNAs targeting different sites within the locus could further reduce the likelihood of resistance allele formation by increasing the frequency of HDR events.

In conclusion, we present an IGD that can efficiently introduce itself and an antiparasitic effector into targeted *A. gambiae* populations. The drive element shows no enhanced vector competence for the most significant pathogens that *A. gambiae* transmits. Despite the observed fitness impacts in nanos^d^ mosquitoes, their shortened lifespan may further aid in reducing transmission. Further studies are required to test this gene drive under more realistic environmental conditions and in combination with other antimalarial effectors.

## Materials & Methods

### Generation of transgenic mosquitoes

The annotated DNA sequence file of the plasmid pD-Cas9-nanos is provided. Briefly, the intron containing the marker module and gRNA module is based on the first *A. gambiae* gambicin 1 intron, which is 111 bp long. Three potential insertion sites for the two modules were tested with the *Drosophila* splice predictor, and the modules were placed between the bases ATA and GGA^39^. The intron was inserted into the E2A autocleavage peptide between the bases TAC and GCC, and the last C was silently recoded to G, to match the endogenous flanking regions in gambicin exons 1 and exon 2. Two fragments were synthesised by GeneWiz Ltd., the first spanning the first part of E2A and the intron, as well as loxP and the rabbit globin terminator 3’ UTR to terminate GFP, and the second one spanning the second part of the intron and E2A. They were cloned into an intermediate plasmid to assemble the entire E2A containing the intron with the GFP module and the *Anopheles gambiae* U6 promoter and scaffold for the gRNA as previously published^40^. Approximately 800 bp of homology arms were amplified from *A. gambiae* Ifakara genomic DNA and incorporated at the 5’ and 3’ ends of the construct. The *nanos* gRNA was chosen using CRISPOR^41^, checked for conservation with the Ag1000G database^42^ and inserted via primers with overlaps during the final Gibson assembly. *A. gambiae* Ifakara eggs were injected with the donor plasmid pD-Cas9-nanos and the Cas9 helper plasmid p155^43^ at a concentration of 200 ng/µL each, yielding 34 G_0_ transients. Five G_1_ transgenics were obtained from a cage with 10 G0 females crossed to wild-type males. The insertion of G_1_ transgenics were confirmed by Sanger sequencing. The marker-free nanos^d^ line was then established by crossing nanos^D^ to a Cre recombinase-expressing line (Eric Marois, unpublished). This resulted in excision of the GFP marker located between two loxP sites. The marker-negative individuals were intercrossed and confirmed to be nanos^d^ homozygous by pupal case PCR.

### Mosquito husbandry

*Anopheles gambiae* Ifakara strains were maintained at ∼27°C and ∼70% humidity with a 12h:12h light:dark cycle and provided with 10% fructose *ad libitum*. All experiments were performed with cow blood (First Link (UK) Ltd.) except for parasite and virus infection experiments (human blood; Cambridge Bioscience).

### Assessment of gene drive

For the *nanos* drive autonomous homing assay, each replicate of heterozygous nanos^D^/nanos^d^ males or females was crossed to wildtypes (pool sizes of 35– 105 individuals), and their progeny were collected for subsequent genotyping. The inheritance of nanos^D^ was determined by the presence of GFP, whereas the inheritance of nanos^d^ was obtained by genotypic PCR with primers 663, 716, and 717. For non-autonomous homing of MM-CP, 45–50 of nanos^d^;MM-CP double-heterozygous males or females were crossed to wild type mosquitoes, the progeny were collected and underwent genotypic PCR for nanos^d^ (using primers 663, 716, and 717) and for MM-CP (using primers 260, 531, and 532). Estimated means and 95% confidence intervals were calculated by a generalised linear mixed model, with a binomial (‘logit’ link) error distribution fitted using the glmmTMB package^44^. Replicates were included as a random effect, and the results were averaged over the levels of sex when comparing the different drives (nanos^D^ and nanos^d^) or loci (*nanos* and *zinc carboxypeptidase A1*).

### Amplicon sequencing

Genomic DNA was extracted with the DNeasy Blood & Tissue Kit (Qiagen) from approximately 400 L2-L3 larvae progeny from male or female heterozygotes nanos^d^ that were crossed to wild-type mosquitoes. PCRs were performed with Q5 High-Fidelity DNA Polymerase (NEB) using primers 717 and 737. Annealing temperature, extension time, and cycle number were set to 63°C, 10s, and 30 cycles, respectively. Amplicons were purified with the QIAquick PCR Purification Kit (Qiagen) and submitted to Amplicon-EZ NGS (GeneWiz Ltd.), and the data were analysed with CRISPRESSO and visualised with R (R Development Core Team).

### Pupation timing and adult survival assays

Three replicates of 100–105 homozygous nanos^d^ or wild-type larvae were maintained in trays under laboratory conditions. The number of days required to develop from the L1 larvae to pupae was recorded, and the sex ratio was determined by pupal sexing. For the adult survival assays, two replicates of 50 nanos^d^ or wild-type adults were maintained separately in male and female cages and provided with 10% fructose under laboratory conditions. The number of dead mosquitoes was recorded daily to monitor the life span of each population.

### RNAseq analysis

Ovaries from 2–4 days post-emergence, unmated homozygous nanos^d^ and wild-type females were dissected. Ovary samples were homogenised with a sterile pestle, and RNA extractions were performed using the Direct-zol RNA Microprep kit (Zymo Research), including the DNase treatment step. A total of 30 ovary pairs per line were collected for three biological replicates. The mRNA samples were purified using poly-T oligo-attached magnetic beads, and cDNA libraries were generated by using dUTP. The cDNA libraires were sequenced on a NovaSeq X Plus Illumina platform (instrument: A01426), generating 150 bp paired-end reads (NAR accession: to be confirmed). Sequencing reads were aligned to the *Anopheles gambiae* PEST genome (AgamP4.13, GCA_000005575.2), supplemented with the nanos^d^ sequence reference, using HISAT2 v2.0.5 with the parameters –dta and –phred33^45^. Differential gene expression was analysed using DESeq2 v1.20.0, and gene ontology (GO) enrichment analysis was conducted using g:Profiler, applying a p-value cut-off of 0.01.

### Parasite and virus infections

Wild-type or nanos^d^ adult mosquitoes were exposed to an infectious blood meal containing mature *P. falciparum* NF54 gametocytes (approximately 0.5% parasitaemia in the infectious blood) using a streamlined standard membrane feeding assay or with *P. berghei* ANKA 2.34 by direct feeding on infected mice (2–3% parasitaemia). Engorged mosquitoes were maintained at 27°C or 21°C for *P. falciparum* and *P. berghei* infections, respectively. For both *P. falciparum* and *P. berghei* infection experiments, 10% fructose was provided to the mosquitoes from 48 hours post-blood meal to eliminate unfed individuals. O’nyong nyong virus Ahero strain (KX771232.1) was used for virus infection experiments. Three to four-day-old adult mosquitoes were provided a mixture of ONNV and human red blood cells at a final titer of 1×10^7^ PFU/mL. The infectious blood feeding was conducted at 37°C for no more than 30 minutes, the mosquitoes were immobilised on ice to remove non-engorged mosquitoes. The engorged mosquitoes were kept at 27°C and 80% relative humidity until salivation. At 7 days post-infection, the mosquitoes were immobilised on ice and their legs and wings were removed for salivation. Saliva was collected by inserting the proboscis into a P20 tip filled with 5 µL of fetal bovine serum (FBS). After 30 min, the saliva-containing FBS was expelled into 45 µL of Dulbecco’s Modified Eagle media and stored at - 80 °C until analysis. After salivation, the mosquitoes’ abdomens and head-thoraxes were separated and stored individually for viral infection and dissemination assays, respectively.

### Population cage experiments

Male and female pupae of nanos^d^ and MM-CP (which carries a *zinc carboxypeptidase A1*-targeting gRNA^31^) mosquitoes were allowed to emerge in separate cages and mixed to mate as adults. For each replicate, 100 heterozygous nanos^d^, 100 heterozygous MM-CP mosquitoes for each sex were seeded at G0, totalling 400 individuals. For each generation of the cage trials, adults were blood fed at approximately six days post-emergence and allowed to oviposit into two egg bowls filled with water and lined with filter paper. The hatched larvae were transferred to trays with 105 larvae each and were maintained until the pupal stage to seed the next generation of the cage trial. For the four replicate cages that include nanos^d^ and the MM-CP transgene, larvae at L2-L3 stage were randomly selected from each generation and placed in a 96-well plate (92 larvae and 4 controls: three DNA extraction buffer and one water controls) and gDNA was extracted with the Phire Tissue Direct PCR Kit (Thermo Scientific). Multiplex PCRs were performed with primers 663, 716, and 717 for nanos^d^, and primers 260, 531, and 532 for MM-CP.

### Statistics, visualisation, and reproducibility

To quantify gene drive inheritance, we analysed data from 15 independent crosses of *A. gambiae* mosquitoes carrying a gene drive under control of the *nanos* promoter. For each cross, the number of *nanos*-IGD (nanos^d^ or nanos^D^) and wild-type F2 progeny was recorded. The resulting dataset consisted of one row per cross and a binomial response variable (*nanos*-IGD^+^, *nanos*-IDG^-^), with the following categorical predictors: F1 genotype at the drive locus, F1 drive carrier sex, and F0 drive carrier sex. A random intercept for cross ID was included to account for overdispersion and between-cross heterogeneity (variance = 0.79, SD = 0.89). Data analysis and visualisation were performed with R. Code to reproduce plotting and statistical analysis performed in R is provided. Gene diagrams were generated with Inkscape.

## Author contributions

Conceptualization: N.W., A.H., Data curation: P.S.Y., S.V., Formal analysis: S.V., P.S.Y., Funding acquisition: N.W., G.K.C., Investigation: P.S.Y., P.C., G.D.C., M.G.I., I.A., M.A.K., Methodology: G.D.C., P.C., P.S.Y., M.G.I., Project administration: N.W., G.K.C., Resources: G.D.C., P.C., M.G.I., P.S.Y., A.H., I.A., M.A.K., Software: S.V., Supervision: N.W., D.V., G.K.C. Validation, P.S.Y., S.V.: Visualization: S.V., Writing – original draft: P.S.Y., S.V., Writing – review & editing: N.W., S.V., P.S.Y., G.K.C.

## Acknowledgments

The authors would like to thank Calvin Yee Kim Guan, Chiamaka Valerie Ukegbu, Temesgen Memberu Kebede and Julia Cai.

## Supplementary figures

**Figure S1.**
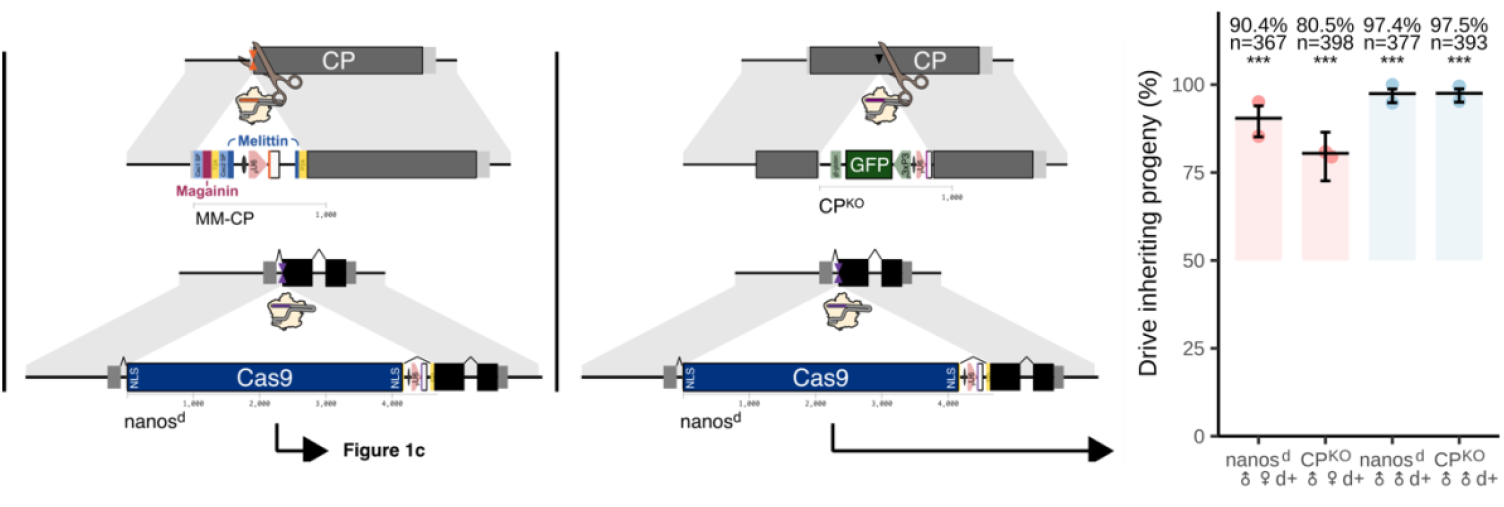
Targeted modifications at the CP locus and homing rates in *nanos*-IGD mosquitoes. Schematic representation of two transgenic integrations at the CP locus and the inheritance rate of nanos^d^ mosquitoes that crossed to CP-KO mosquitoes. (Left) The MM-CP construct incorporates magainin-2 and melittin under the regulation of CP promoter. (Middle) The CP-KO construct includes a GFP marker, disrupting CP expression. (Right) Inheritance rates of nanos^d^ and CP-KO mosquitoes, measured as the percentage of progeny inheriting nanos^d^ and CP-KO. Statistical significance is indicated by asterisks (*).

**Figure S2.**
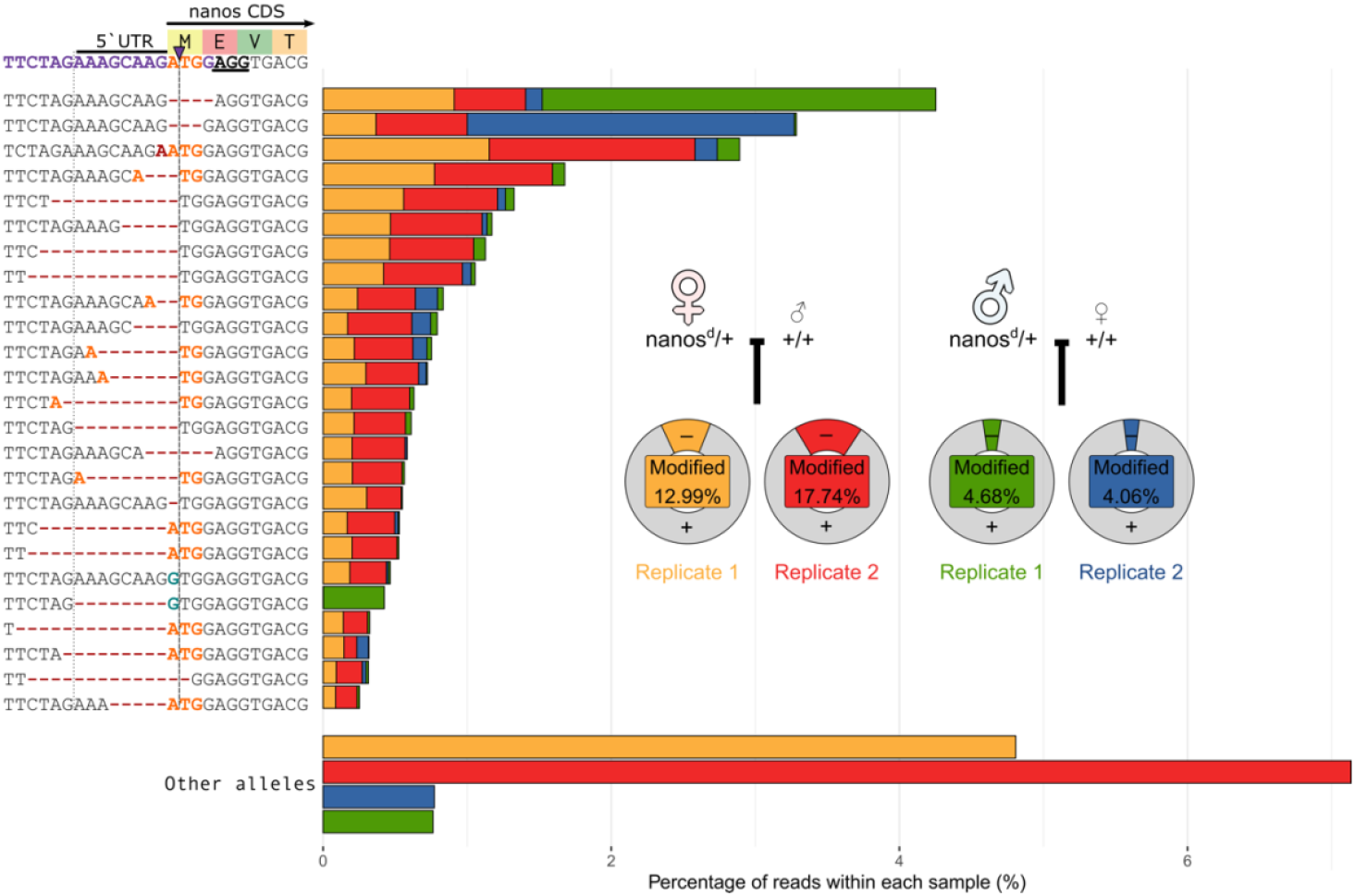
Top 25 most abundant nanos^d^ resistance alleles found from male and female crosses. Mutant alleles that recreate an ATG start codon are highlighted. Blue and green bars represent the percentage of each genotype from the two male crosses, whereas red and orange bars represent the two female crosses.

**Figure S3.**
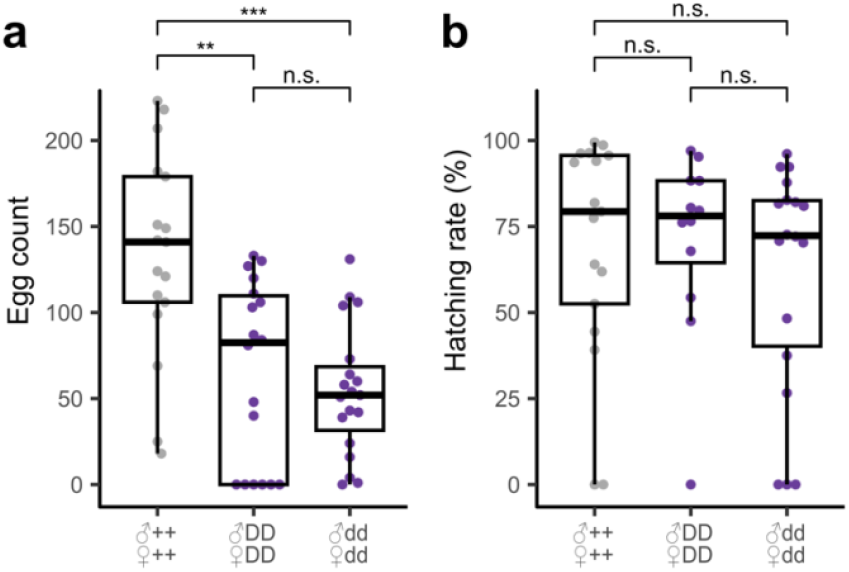
Fecundity and fertility of homozygous nanos integral gene drive mosquitoes with and without the intronic GFP cassette. The number of eggs **(a)** and hatching rate **(b)** were recorded for intercrossed nanos^D^ or nanos^d^ females. DD indicates homozygous nanos^D^; dd indicates homozygous nanos^d^. Egg count significance levels were calculated using an unpaired two-tailed Student’s *t*-test using Bonferroni correction. Hatching rate means and significance levels were calculated using a binomial GLM with replicate as a random effect.

**Figure S4.**
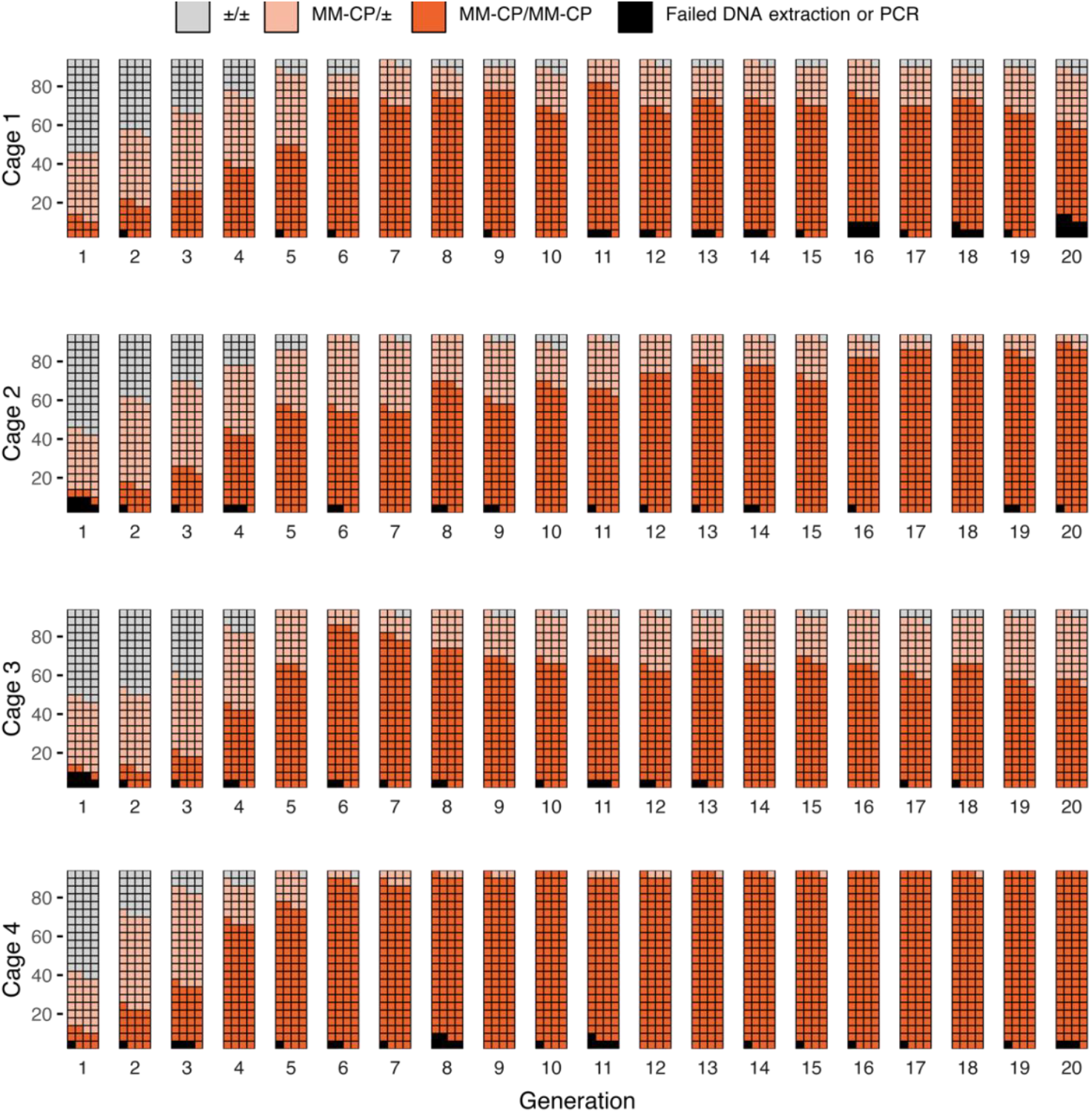
Construct and inheritance patterns of the MM-CP during the cage trials. Schematics represent the composition of MM-CP alleles across 20 generations in the four cage populations, each unit represents an individual mosquito.

**Figure S5.**
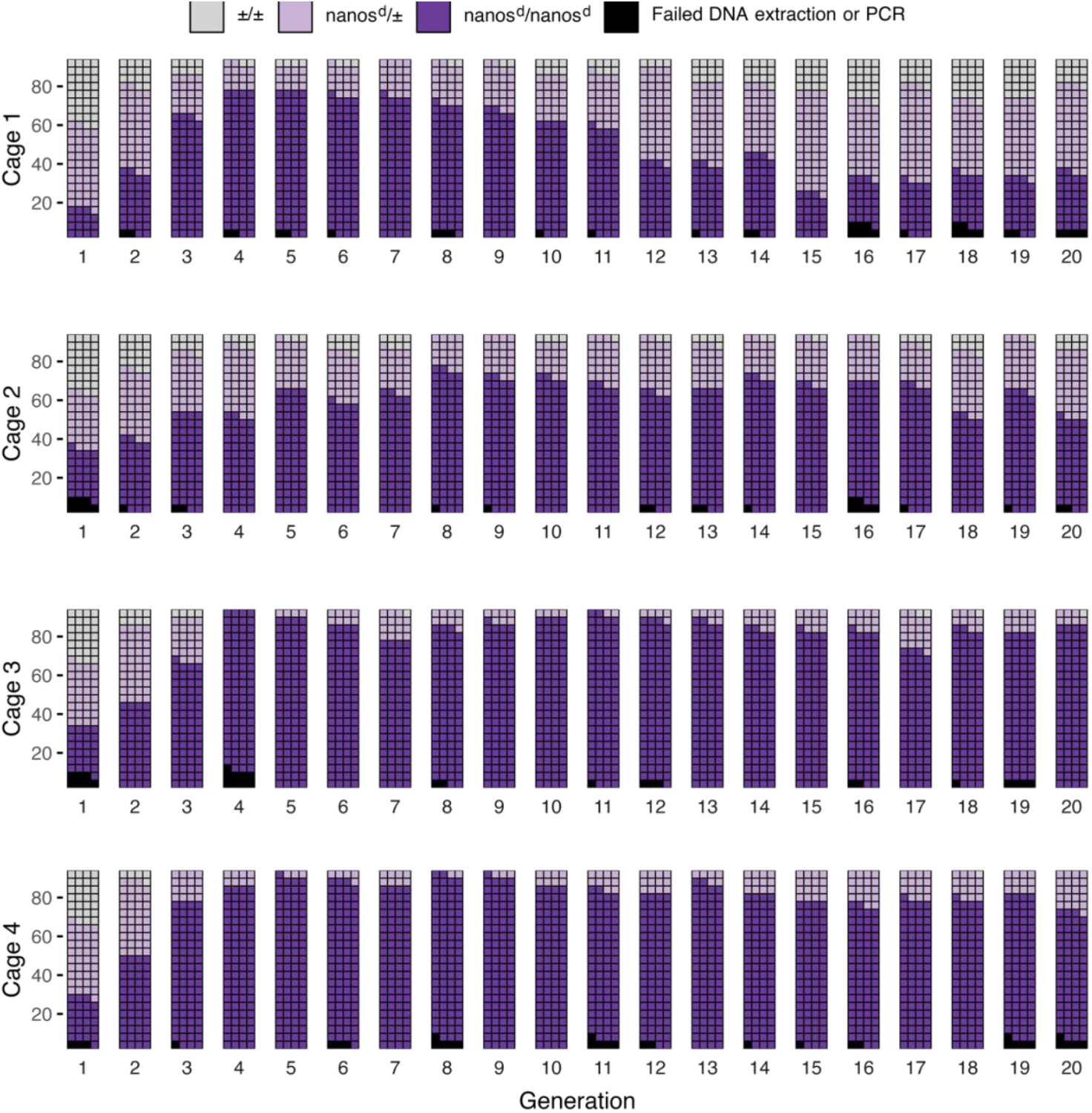
Inheritance patterns of the nanos^d^ during the cage trials. Schematics represent the composition of nanos^d^ alleles across 20 generations in the four cage populations, each unit represents an individual mosquito.

